# Structural and thermodynamic analyses of human TMED1 (p24γ1) Golgi dynamics

**DOI:** 10.1101/2021.02.13.431076

**Authors:** Danielly C. A. M. Mota, Renan M. Mori, Mariana R. B. Batista, Luis G. M. Basso, Iara A. Cardoso, M. Cristina Nonato, Antonio J. Costa-Filho, Luis F. S. Mendes

## Abstract

The transmembrane emp24 domain-containing proteins (TMED), also called p24 proteins, are members of a family of sorting receptors present in all representatives of the domain Eukarya and abundantly present in all subcompartments of the early secretory pathway, namely the endoplasmic reticulum (ER), the Golgi, and the intermediate compartment. Although essential during the bidirectional transport between the ER and the Golgi, there is still a lack of information regarding the TMEDs structure, oligomerization propensity, and biophysics of their interactions with the transport cargo. Here, we describe the first high-resolution structure of the Golgi dynamics (GOLD) domain of a TMED1 representative and its biophysical characterization in solution. The crystal structure showed a dimer formation that is also present in solution in a salt-dependent manner, suggesting that the GOLD domain can form homodimers even in the absence of the TMED1 coiled-coil region. A molecular dynamics description of the dimer stabilization, with a phylogenetic analysis of the residues important for the oligomerization and a model for the orientation towards the lipid membrane are also presented.

## 1. Introduction

The transport of protein cargo in the heart of the secretory pathway is mediated by coated vesicles and multiple accessory proteins [1-4]. In the early secretory pathway, the transport of proteins between the endoplasmic reticulum (ER) and the *cis*-Golgi face (anterograde transport) is mediated by the well-known coat protein complex II (COPII) [5], whereas the retrograde transport is controlled by the coat protein complex I (COPI) [6]. The COPII machinery selects the well-folded and active cargo at the ER exit sites for further export in membrane-bound vesicles toward the Golgi [7,8]. To correctly deal with the great number of transport cargos, which can reach up to 15% of the total proteome in humans [9], protein sorting receptors are required in great number and diversity [3]. These sorting receptors are transmembrane proteins that recognize complex carbohydrates and/or polypeptide signals in the transport cargo, and that are constantly cycling between the ER and the Golgi in vesicle intermediates formed by COPI and COPII [3].

One sorting receptor family present in all representatives of the domain Eukarya is the transmembrane emp24 domain-containing protein (TMED), also called p24 family [10-12]. Members of this family have an average molecular mass of 24 kDa and their number varies for each species [12]. The TMED proteins emerged as central regulators in the secretion of biomolecules, acting in their cellular traffic and influencing the composition, structure, and function of the ER and the Golgi [12]. Therefore, TMEDs have been associated with the progress of various human diseases, such as Alzheimer’s disease [13] and carcinomas [14,15], as well as have shown specific roles in the cell signalling process [12], insulin secretion [16], and cell growth [17].

Hitherto, ten representatives have been found in mammalians [18], including one member (TMED10) whose allele disruption was shown to lead to early embryonic lethality [19]. The human genome is an exception encoding for only nine TMEDs because the TMED11 gene contains an in-frame stop codon, although it has been observed in other primates [20]. TMEDs are divided into four subfamilies based on sequence identity (α, β, γ, and δ) [21]. Albeit their low sequence identity, even within the same subfamily, all TMEDs are predicted to share a similar domain organization [12]: a small cytosolic portion at the C-terminus with a motif for interaction with the COPs, a single transmembrane domain, a coiled-coil domain, a flanking region, and a long luminal region comprising a domain known as Golgi Dynamics (GOLD) [12].

Currently, structural data are available only for the isolated GOLD domains from mammalian TMED2, TMED5, and TMED10 [22,23], which showed a structural organization of a β-sandwich fold composed of two four-strand antiparallel β-sheets connected by a conserved disulphide bond [22,23]. There are “visually” two different “sides” in this conformational organization, called concave (formed by β2, β7, β4, and β5) and convex sides (formed by β6, β3, β8, and β1) [22,23]. The known 3D structures of the GOLD domains span three of the four TMED subfamilies: TMED2 belongs to the subfamily β, while TMED5 and TMED10 belong to the subfamilies γ and δ, respectively [12].

*In vivo* studies suggested that TMED1, one of the TMED family members, is involved in interleukin-33 (IL-33) signalling by mediating the transport of the ST2 receptor (member of the Toll-like/IL-1-receptor superfamily) to the cell membrane [24]. IL-33 mitigates sepsis by increasing the number of neutrophils at the site of infection, thus increasing the clearance of bacteria [25]. The ST2 receptor not only interacts with IL-33, generating T-cell-mediated immune responses, but is also involved in infectious diseases, asthma, and allergic responses [26,27]. The pair IL-33/ST2 is also implicated in cardiovascular diseases [28], which suggests that the IL33/ST2 signalling could be not only a cardiovascular biomarker but also a possible therapeutical target for the prevention of heart failure [28]. In IL-33 signalling, the ST2/TMED interaction involves the TIR domain of the ST2 receptor and the GOLD domain of TMED1 (tp24 or p24γ1) [24]. Interestingly, TMED1 plays a role in the expression of other interleukins, such as interleukin-8 and interleukin-6 [24]. Therefore, the GOLD domain of TMED members are, at least indirectly, part of the signalling processes for the immune system and cell differentiation [29].

To expand the limited structural knowledge of TMED proteins, we performed biophysical analyses of the GOLD from the human T1/ST2 receptor-binding protein TMED1 (UniProtKB - Q13445). We also provide the first structural and thermodynamic characterization of a TMED1 representative, shedding light on the structural organization, stability, and oligomerization propensity of the GOLD domains within the TMED family.

## 2. Materials and Methods

### 2.1 Protein expression and purification

The human TMED1 gene was synthesized with codon optimization for *E. coli* expression by GenScript^®^ Biotech. The signal peptide-free GOLD domain of TMED1 was amplified from the TMED1 gene from the amino acid residues 24 to 130, plus a Tobacco Etch Virus (TEV) site at its N-terminus and subcloned in a pET28a vector using the Nde1 and Xho1 restriction sites. Protein expression was performed in Origami™ 2 (DE3) competent cells (Novagen) using LB medium. Briefly, cell solution was allowed to grow to an OD of 0.8 at 37°C and under 200 rpm agitation, and then induced by using 0.5 mM of isopropyl-β-D-thiogalactoside (Sigma Aldrich), followed by incubation for 18 hours at 18°C and 200 rpm. Cells were isolated by centrifugation in a Sorvall RC6+ (*ThermoScientific)* per 10 minutes at 7,000xg, resuspended in a PBS (Phosphate Buffered Saline) (1X, pH 7.4) + 0.5% Triton X-100 solution and disrupted by sonication in a Branson 450 Digital Sonifier® (Sonitech).

The soluble fraction of the disrupted cell solution was isolated by centrifugation at 14,000xg for 20 minutes and loaded into a Ni-NTA column system (Qiagen). The column was washed with a volume corresponding to 5 column volumes (CV) of 1X PBS and 5 CV of 1X PBS + 20 mM imidazole. The bound protein was eluted using 3 CVs of 1X PBS + 500 mM imidazole. The protein solution was then concentrated using a 3 kDa cutoff centrifugal filter unit (Millipore, Burlington, MA, USA) and loaded into a Mono Q 5/50 GL (GE Healthcare Life Sciences, Chicago, IL, USA) coupled to an Äkta purifier system (GE Healthcare). The protein solution was then submitted to a linear gradient of NaCl from 20 to 500 mM (+ 20 mM Tris-HCl, pH 8.0) in a flow rate of 0.5 mL/minute with a 1 mL volume of injection. Protein concentration was estimated using the theoretical extinction coefficient of 6.085 M^-1^ .cm^-1^ at 280 nm. The success of protein purification was accessed by using 15% SDS-PAGE in a Bio-Rad Mini-PROTEAN® Tetra Vertical Electrophoresis Cell.

### 2.2 Circular Dichroism (CD)

Far UV-CD spectra (195-260 nm) were measured in a Jasco J-810 spectrometer (Jasco Corporation, Japan) equipped with a Peltier control system and using a quartz cell with a 1 mm pathlength. The spectra were recorded from 260 to 195 nm, with a scanning speed of 100 nm/min, spectral bandwidth of 1 nm, and response time of 0.5 s. All the protein stock solutions were at a minimum concentration where the dilution in a 20 mM sodium phosphate pH 8.0 was at least 20-fold, in a final protein concentration of 0.15 mg/mL. Spectral deconvolution was performed using BeStSel [30]. Thermal denaturation was performed by monitoring the ellipticity at 205 nm in the range from 20 to 80°C using a heating rate of 60°C/h.

### 2.3 Size Exclusion Chromatography with Multi-Angle Light Scattering (SEC-MALS)

SEC-MALS measurements were performed on a miniDAWN TREOS multi-angle light scattering equipment with detectors at three angles (43.6°, 90° and 136.4°) and a 659 nm laser beam (Wyatt Technology, CA). A Wyatt QELS dynamic light scattering module for determination of hydrodynamic radius and an Optilab® rEX refractometer (Wyatt Technology) were used in-line with a size exclusion chromatography analytical column (Superdex 75 HR10/300, GE Healthcare). BSA was used as a control. The protein solutions (∼2 mg/mL) were eluted in a 50 mM Tris HCl, 500 mM NaCl buffer, pH 8.0 with a flow rate of 0.5 mL/min. The data were processed using ASTRA7 software (Wyatt Technology) with the following parameters: refractive index of 1.331, 0.890 cP for the viscosity of the solvent and a refractive index increment of 0.1850 mL/g. Protein solutions were centrifuged for 10 minutes at 10,000xg at 4°C prior to use.

### 2.4 Size Exclusion Chromatography

Size-exclusion chromatography was carried out using a Superdex 75 (HR 10/30) coupled to an Äkta purifier system (GE Healthcare), equilibrated with a 50 mM Tris/HCl, pH 8.0 and varying the NaCl concentrations (20 and 500 mM). Apparent molecular mass was obtained from plots of K vs. log (MM) (molecular mass), where K is the partition coefficient. Standard proteins used were from the gel filtration calibration kit LMW and HMW (GE Healthcare) with the exception of *Cryptococcus neoformans* acyl-CoA binding protein (ACBP) that was purified as reported in the literature [31]. TMED1 GOLD domain oligomerization analyses, together with the standard proteins, were performed in 50 mM Tris/HCl pH 8.0 + 20 mM NaCl using a 100 µL loop injection and a flow rate of 0.5 mL/minute.

### 2.5 Differential Scanning Calorimetry (DSC)

DSC measurements were carried out with the protein solution eluted from the ionic exchange chromatography and after dialysis in 20 mM Tris-HCl, pH 8.0 with 20 or 300 mM NaCl. The experiments were performed on a VP-DSC MicroCal MicroCalorimeter (Microcal, Northampton, MA, USA) using a heating rate of 1°C/min. The thermograms were recorded from 20 to 95°C, at a controlled pressure of 1.6 atm. Instrumental buffer baselines were recorded prior to the protein unfolding experiments to register the thermal history of the calorimeter. The raw DSC traces were subtracted with the buffer baseline and then normalized by protein concentration (130 µM). Three different baselines (cubic, progress, and manual) were reconstructed using the Microcal Origin software and subtracted from the normalized thermograms. Simulations of the thermograms were carried out using CalFitter v.1.3 [33]. All experiments were repeated at least twice.

### 2.6 Crystallization, data collection and structure resolution

Crystallization experiments were carried out by the sitting drop vapor diffusion method, using 10 mg/ml of protein concentration at 21°C. Drops were prepared by mixing 1.0 μL of protein solution and 1.0 μL of reservoir solution. Crystals of TMED1 GOLD domain were obtained in the presence of 0.1 M sodium acetate trihydrate pH 4.6 + 8% w/v PEG 4,000 (Crystal Screen 1 #37, Hampton Research), and started to appear after approximately 14 days of incubation. The crystals were soaked in a cryoprotectant solution (sodium acetate pH 4.5; 30% 2-methyl-2,4-pentanediol, MPD) prior to cooling in liquid nitrogen. The data set was collected at - 173°C on the PROXIMA 1 protein crystallography beamline at SOLEIL, France, using an EIGER-X 16M detector (Dectris, Baden, Switzerland). 3,600 frames with an oscillation step of 0.1° were collected using an exposure time of 0.1 s per image with a crystal-to-detector distance of 224 mm. X-ray diffraction data were processed and scaled with *XDS* [34] and *AIMLESS* [35]. The structure was solved by molecular replacement using the Molrep [36] contained within CCP4 software suite [35]. The coordinates of murine TMED5 (p24γ2) GOLD domain (PDB ID 5GU5) were used as template. Automated model rebuilding was performed using Buccaneer [37], and the structure was refined with REFMAC5 [38], followed by manual map inspection and model building using Coot [39]. The quality of the model was regularly checked using MOLPROBITY [40]. The figures were prepared using CCP4MG [41]. The residues in the PDB file were numbered considering all the non-native amino acids present in the N-terminus coming from the pET28a expression vector. The amino acid sequence is presented in the supplementary material and this numbering is also the one used in the manuscript.

### 2.7 Phylogenetic analyses

Sequences of the GOLD domain of the TMED γ subfamily were collected based on their similarities with the sequence used for TMED5 GOLD domain crystallization and the signal peptide in the N-terminus for each sequence was predicted using SignalP-5.0 Server [42]. The evolutionary history was inferred by using the Maximum Likelihood method and JTT matrix-based model [43]. The tree with the highest log likelihood (−1.071,52) is then shown. Initial tree(s) for the heuristic search were obtained automatically by applying Neighbor-Join and BioNJ algorithms to a matrix of pairwise distances estimated using a JTT model, and then selecting the topology with superior log likelihood value. The tree is drawn to scale with branch lengths measured in the number of substitutions per site. Evolutionary analyses were conducted in MEGA X [44]. Protein sequences were aligned using Clustal Omega [45] and the figure, with the statistical analyses, was generated using Jalview (https://www.jalview.org/ accessed in 2020). There is a shift of 28 between the numbered residues in the sequence alignment and the numbered residues in the PDB file. This is due to the non-native amino acids carried in the protein expression as a tag in the N-terminus.

### 2.8 Molecular dynamics simulations

Initial coordinates for the TMED1 GOLD monomer and dimer used in the simulations were obtained from the crystallographic structure here reported and refined to 1.7 Å resolution. A solvation shell of at least 15 Å was created using PACKMOL [46,47], and sodium and chloride ions added in a concentration close to 0.15 mol L^-1^ to render the system electrically neutral. All simulations were performed with NAMD [48] using periodic boundary conditions and CHARMM36 [49] parameters for proteins and TIP3P [50] for water. A 12 Å cut-off radius was used for the van der Waals interactions, whereas the electrostatic forces were handled by Particle Mesh Ewald sums [51]. Simulations were performed at 298.15 K and 1 atm using a time-step of 2 fs. Temperature was controlled using Langevin dynamics with a friction coefficient of 10 ps^-1.^ The Nosè-Hoover algorithm was used for pressure control with a piston oscillation period of 200 fs and a decay rate of 100 fs. Covalent bonds involving hydrogen atoms in the protein were constrained to their equilibrium distances using the SHAKE algorithm, while SETTLE [52] was used for water. The solvated structures were subjected to the following steps of minimization and equilibration: 1) 1,000 CG (conjugate gradient) energy minimization steps and 200 ps MD keeping the position of all residues fixed; 2) 500 CG energy minimization steps and 200 ps MD keeping all Cα carbons fixed; 3) 2 ns MD with the entire system allowed to move. From the equilibrated systems, 3 independent 100 ns simulations for each system were carried out. The surface area buried at the dimer interface was calculated as the sum of the solvent accessible area of the monomers minus the solvent accessible surface area of the complex, divided by two. The calculations were performed with VMD with water represented as a sphere of radius 1.4 Å [53,54].

### 2.9 MM/PBSA free energy calculation

The calculation of binding free energy, ΔG_bind_, between monomer A and monomer B to form a dimer complex were evaluated using MM/PBSA (Molecular Mechanics-Poisson Boltzmann Surface Area) method [55,56] as implemented in AMBER18 [57,58]. In the MM/PBSA method, the dimerization free energy (ΔG_bind_) between each monomer to form the dimer complex was calculated as:

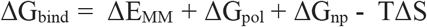

where ΔE_MM_, ΔG_pol_, and ΔG_np_ represent the gas-phase molecular mechanical energy change, the polar contribution to the solvation free energy, and the non-polar contribution, respectively. In MM/PBSA method, the polar solvation term was calculated by solving the linearized Poisson-Boltzmann (PB) equation and the non-polar term was determined based on the solvent accessible surface area (SASA). The term TΔS represents the conformational entropy, which can be estimated by normal-mode analysis. All MM/PBSA analysis were carried out using the MMPBSA.py [59] python script in the AmberTools18 [57]. The calculations were performed on 600 spaced snapshots extracted from the production MD runs (200 snapshots for each production MD run). The salt molar concentration was set to 0.1 M in solution. The trajectory and structure files from NAMD were transformed into Amber compatible file via *cpptraj* and *chamber*, respectively.

## 3. Results and Discussions

### 3.1 TMED1 GOLD heterologous expression and phylogeny

The expression and purification of a GOLD domain from a γ1 representative had no protocol reported hitherto. The heterologous expression of the recombinant protein using standard *E. coli* BL21-derivative lineages showed a very low yield, which could be due to the inherent difficulties of expressing disulphide bond-forming proteins using those lineages. The problem was overcome by using *E. coli* Origami 2 (DE3) cells and the purification from soluble fractions was performed using a two-step purification step (Figure 1A).

**Figure 1:**
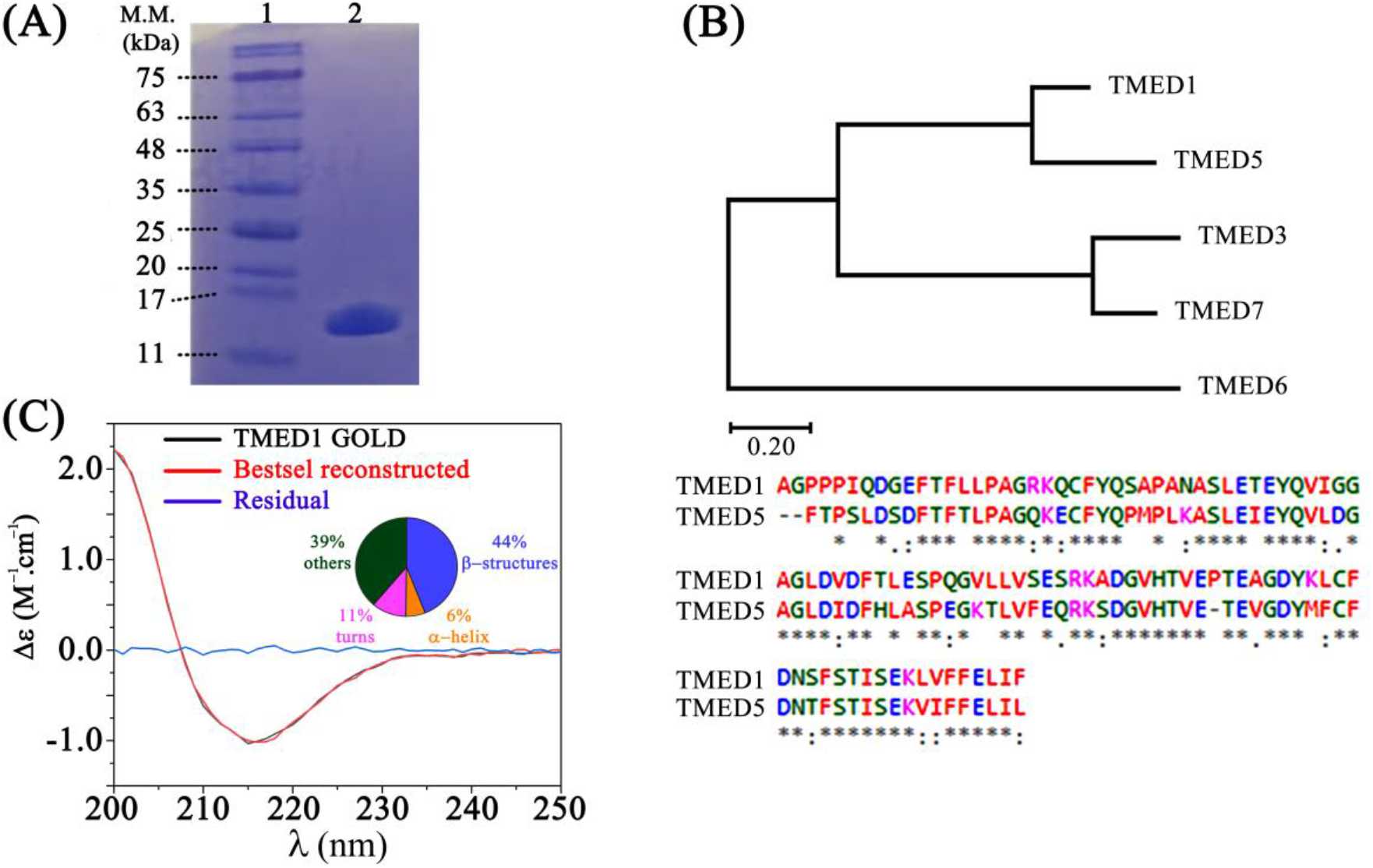
General analyses of TMED1 GOLD domain. (A) SDS-PAGE analysis of the protein purification. In sequence, 1) Molecular mass standard. 2) Final protein solution after the 2-step chromatographic purification. (B) Phylogenetic analyses of the TMED γ subfamily and the sequence alignment between TMED1 and TMED5 GOLD domains. (C) Experimental CD data and theoretically reconstructed spectrum using BeStSel [30]. The inset shows the fraction of each secondary structure element predicted for this protein in solution, with an NRMSD of 0.0217. The figure was built using Adobe Fireworks CS6 and Origin® 8.5.

As a member of the γ subfamily, the TMED1 GOLD domain is closer to TMED5 than to the remaining TMEDs in a phylogenetic perspective (Figure 1B). The sequence identity between the TMED1 and TMED5 is 66% (Figure 1B), being the TMED5 GOLD the only γ representative with known high-resolution structure (PDB ID 5GU5). The TMED1 GOLD domain is well-folded in solution and formed mainly by β-strand structures as inferred from its CD spectrum (Figure 1C), akin to TMED5 GOLD [23].

### 3.2 Crystal structure of the TMED1 GOLD construct shows a highly conserved β-sandwich fold

Up to now, there was no structural characterization of any TMED1 representative. The crystal structure of TMED1 GOLD domain was determined by X-ray diffraction to 1.7 Å resolution. Each asymmetric unit contains three copies of the signal peptide-free GOLD domain comprising residues Ala29 to Asp135 of the human TMED1 (UniProtKB Q13445). In the crystallographic structure, the residues 29 - 32, and 132 - 135 were excluded from the model due to the lack of interpretable electron density. The final structure contains three polypeptide chains (A, B, and C), and 94 solvent sites treated as water oxygens. The final round of refinement reached R_work_ of 21.2% and R_free_ of 23.6%. The crystallographic data were summarized in **Table 1**. The coordinates and structure factors were deposited in the Protein Data Bank (PDB) with the accession code 7K49.

**Table 1:**
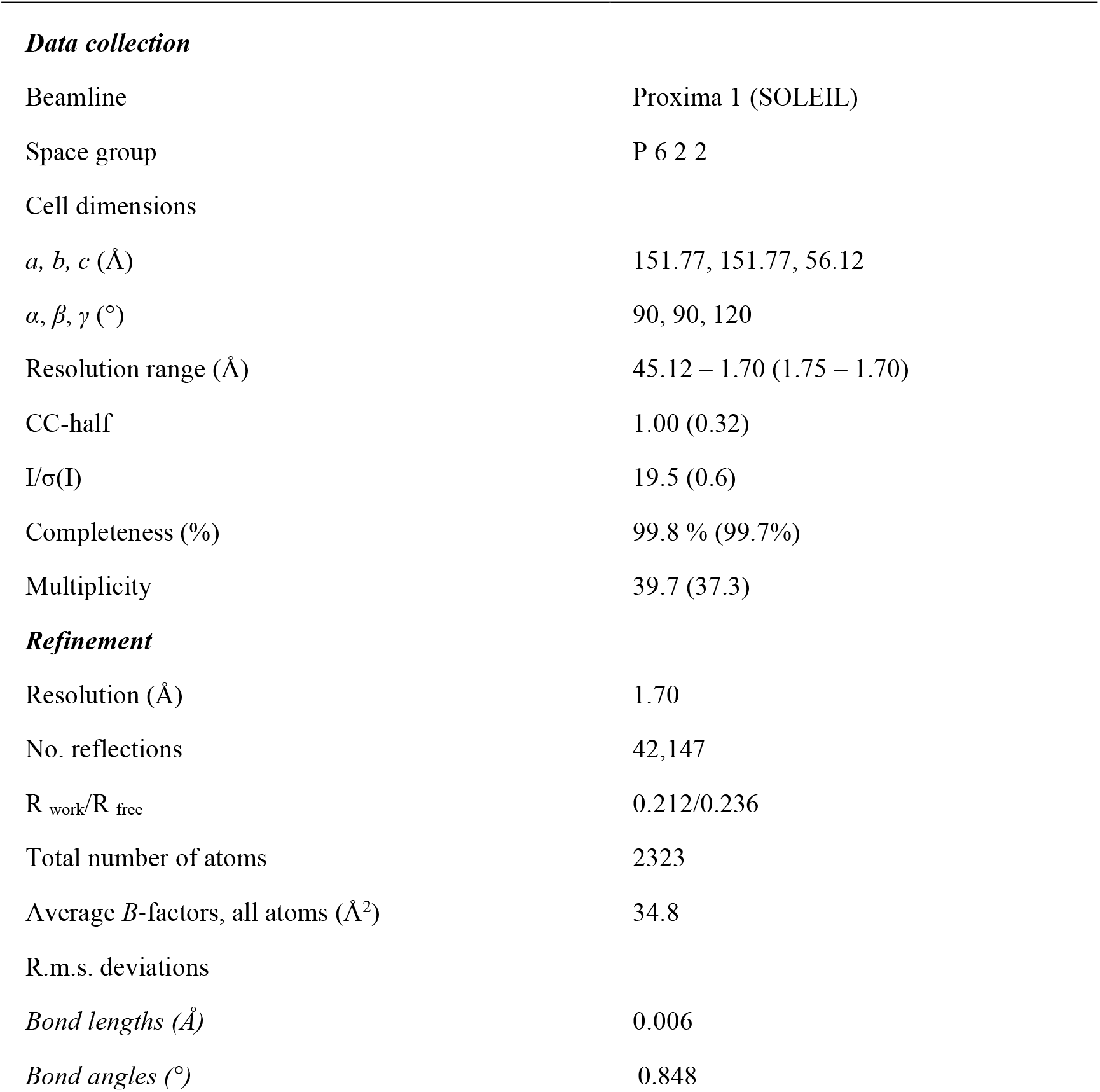
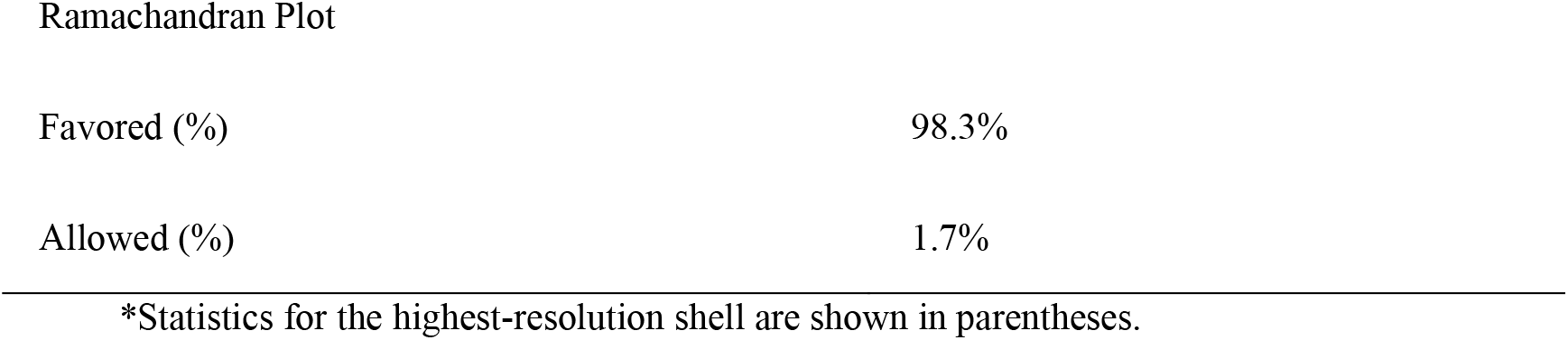
Crystallographic data.

| ***Data collection*** |  |
| --- | --- |
| Beamline | Proxima 1 (SOLEIL) |
| Space group | P 6 2 2 |
| Cell dimensions |  |
| $a, b, c$ (Å) | 151.77, 151.77, 56.12 |
| $\alpha, \beta, \gamma$ (°) | 90, 90, 120 |
| Resolution range (Å) | 45.12 – 1.70 (1.75 – 1.70) |
| CC-half | 1.00 (0.32) |
| $I/\sigma(I)$ | 19.5 (0.6) |
| Completeness (%) | 99.8 % (99.7%) |
| Multiplicity | 39.7 (37.3) |
| ***Refinement*** |  |
| Resolution (Å) | 1.70 |
| No. reflections | 42,147 |
| $R_{\text{work}}/R_{\text{free}}$ | 0.212/0.236 |
| Total number of atoms | 2323 |
| Average $B$-factors, all atoms (Å<sup>2</sup>) | 34.8 |
| R.m.s. deviations |  |
| *Bond lengths* (Å) | 0.006 |
| *Bond angles* (°) | 0.848 |
| Ramachandran Plot |  |
| Favored (%) | 98.3% |
| Allowed (%) | 1.7% |

\*Statistics for the highest-resolution shell are shown in parentheses.

The TMED1 GOLD domain presents the β-sandwich fold seen in other TMED GOLD domains [22,23]. The β-sandwich contains eight β-strands arranged as four strands (β2, β7, β4, and β5) in its concave side and the other four (β6, β3, β8, and β1) in the convex side (Figure 2A). Two cysteine residues, Cys50 and Cys111, form a disulphide bond that bridges β2 and β7 strands, indicating a structural role in the TMED1 GOLD domain fold (Figure 2B). This disulphide bond formed by the two conserved cysteines is also seen in the crystal structures of the TMED 2, 5, and 10 GOLD domains [22,23]. Although conserved and water-exposed, and at least essential for protein stability, hypothesis supported by the failure to express this protein without using the Origami 2 (DE3) strain, it remains unclear whether that disulphide bond could play any functionalities in protein traffic. Unlike the TMED5 GOLD structure (the other member of the γ subfamily), in our case, the short α-helix in the C-terminal end, previously hypothesized as essential for crystallization, is absent (Figure 2C). The structural superposition of the monomeric GOLD domains of human TMED1 and Mus musculus TMED5 results in a C_α_ RMSD of only 1.10 Å (Figure 2C). The superposition of monomeric GOLD TMED1 with monomeric GOLD TMED2 and TMED 10 yields a C_α_ RMSD of 1.20 and 1.64, respectively, suggesting a highly conserved tertiary structure of the GOLD domain among the TMED subfamilies.

**Figure 2:**
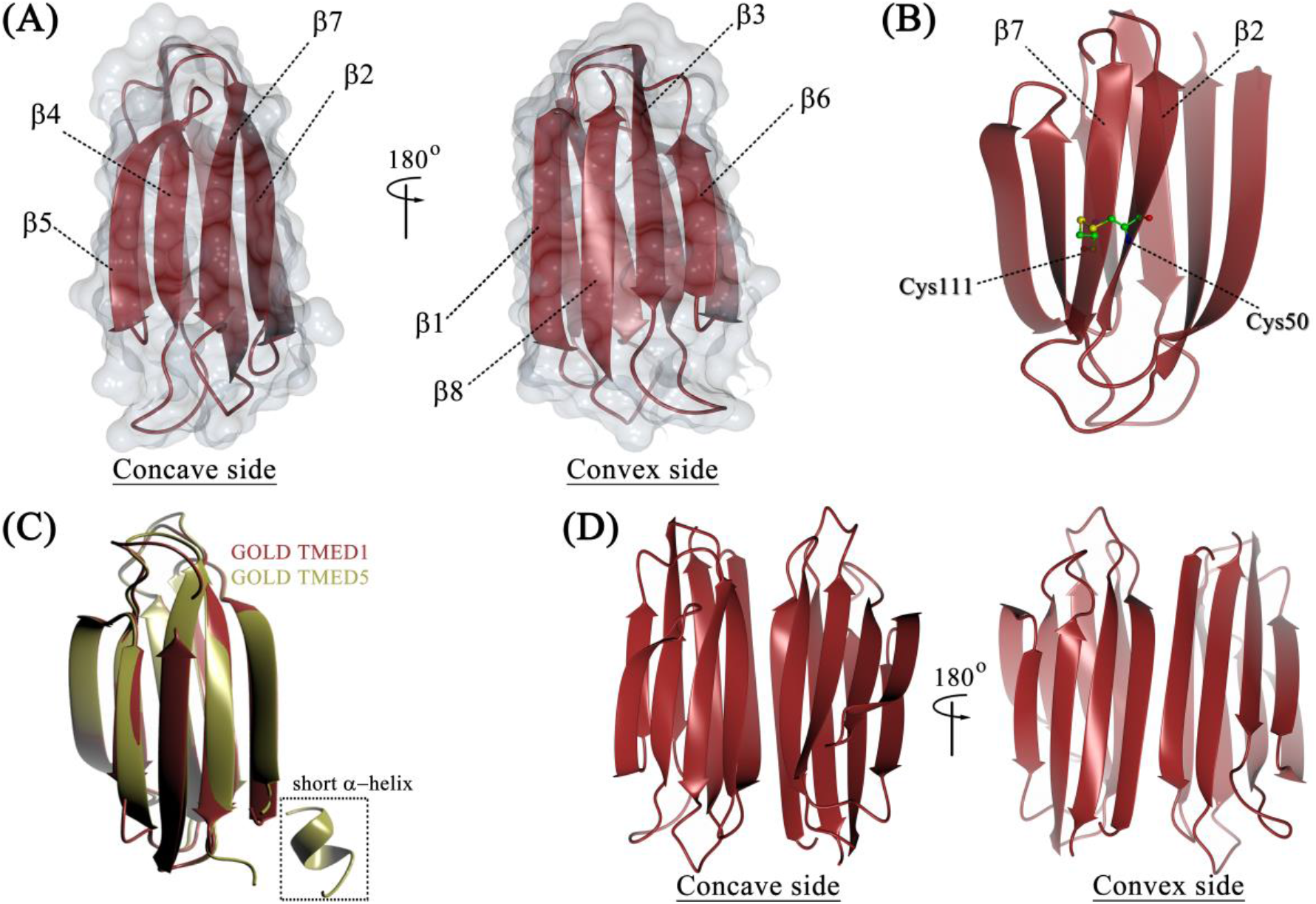
Structural analyses of the TMED1 GOLD. (A) Cartoon representation of the crystallographic structure with a partially transparent surface of both the concave and convex sides. Each β-strand is sequentially labelled. (B) Representation of the disulphide bond formed by both cysteines 50 and 111 connecting the β2 and β7 strands. (C) Structure superposition of the TMED1 (dark red) and TMED5 GOLD (gold) monomers using the Gesamt/SSM method, as implemented in CCP4MG [41]. (D) Cartoon representation of the TMED1 GOLD dimer observed in the asymmetric unit. The dimerization preserves the concave and convex sides observed in the monomer structure. The figure was prepared using Adobe Fireworks^®^ CS6 and CCP4MG.

### 3.3 The GOLD domain forms oligomers in a salt-dependent way

There is an extensive discussion on TMED oligomerization in the literature, both in the homo- and hetero-associations. For instance, TMED2 and TMED10 GOLD domains were observed to form heterodimers [22] suggested as the primary receptor for the Arf1-GDP, the small GTPase involved in COPI-vesicle formation [60]. Hetero-oligomers were also reported for other TMEDs using biochemical approaches, including the first suggestion that TMED would form hetero-tetramers composed of one member of each subfamily [61]. However, further studies showed that TMEDs occur mostly as either monomers or dimers, depending on their subcellular localization [10,62].

Despite the debate regarding its occurrence *in vivo*, TMED oligomerization was intuitively assumed to be driven by the protein’s coiled-coil region [63,64]. Crystallographic analyses of the murine TMED5 GOLD domain suggested the possibility of a dimer formation based on an interaction of one monomer in the asymmetric unity with a symmetry-related molecule using the operator (x, x-y, -z + 2/3) [23]. In this model, the dimer is stabilized by the β1 and β2 strands of each chain and the isolated participation of the C-terminal short helix filling the space between both β2 strands [23]. Although the authors assumed the formation of the dimer, no additional data to contest a hypothesis of a dimerization artifact caused by the crystal packing was provided. In our crystallographic structure, the TMED1 GOLD domain was also found to form a dimeric structure either by applying the symmetry operator (-y,-x,-z) on chain B or within the asymmetric unity, involving the A and C chains (Figure 2D). Akin to the reconstructed dimer of TMED5 GOLD, the interaction between each chain is stabilized by the β1 and β2 strands of each TMED1 GOLD monomer, which are structurally organized in an antiparallel orientation (Figure 2D). Although both dimer models involve the β1 and β2 strands, there is a larger contact area between opposite strands in the TMED1 GOLD structure. The β2 strands interact directly with each other while in TMED5 they are linked via the short α-helix region [23]. Despite the larger contact area involving the β1 and β2 strands observed in TMED1 GOLD dimer, its total dimerization surface area (∼ 763 Å) is smaller than TMED5 GOLD dimerization area (∼ 957 Å). However, the Cα RMSD is approximately 13 Å for their dimeric superposition, indicating a great disparity in the oligomerization arrangement (Figure S1). On the other hand, superposition between the structure of TMED1 GOLD dimer with the proposed TMED10 GOLD dimer (PDB: 5AZY) reveals a Cα RMSD value of approximately 2 Å, despite the low identity between their sequences and the fact that TMED10 is a member of the δ subfamily. Thus, TMED1 and TMED10 dimerize via similar direct interactions involving β1 and β2 strands, and TMED10 dimerization surface area (∼ 815 Å) is also similar to the area observed for the TMED1 GOLD dimer structure.

To get more information on the proposed TMED1 GOLD dimer, the structure was also analysed by MD simulations. To assess the stability of the simulations for the monomer and dimer models, the RMSD values for the backbone of the entire protein along all trajectories were calculated using the crystallographic structure as a reference. The Cα RMSD for both the monomer and the dimer remained relatively low throughout the simulations (Figure S2). The higher RMSD values obtained for the isolated monomer compared to both monomers in the dimer model is likely due to the increased flexibility of the β3/β4-connecting loop (residues 68 to 75) rather than reflecting any significant conformational change or structural destabilization (Figure S2A-B). Moreover, the centre of mass (COM) distance between the monomers was monitored during the simulations (Figure S2C). As shown by the RMSD and the COM distance, the TMED1 dimer is stable during the MD simulations and remains close to the starting crystallographic structure.

We also examined the free energy contributions to the dimer stabilization by performing MM/PBSA calculation of the TMED1 GOLD dimer. The free energy of binding is composed by both an enthalpic and an entropic term. The first includes the contribution from the molecular mechanics and the solvation parts. The second accounts for the differences in the translational, rotational, and vibrational entropies. The TMED1 GOLD dimer is enthalpically stabilized by approximately −28 kcal/mol. For systems with monomers of similar molecular mass, the entropic term is small when compared with the enthalpic term [56,65]. To gain a detailed picture of the dimerization mechanism, the ΔG_bind_ was decomposed on a per-residue basis (Figure 3A). The most significant contributions to the free energy of binding came from the residues Phe39, Thr40, Phe41, Pro44, Arg47, Phe51, Tyr52, Asp107, and Lys122. These residues are located at the dimerization interface and establish important interactions to stabilize the complex. The side chains of the charged residues Asp36, Arg47, Asp107, and Lys122 from β1, β2, β7, and β8 form intermolecular hydrogen bonds (Figure 3B). Another hydrogen bond observed in the simulations involves the main chain of Tyr52 and the side chain of Gln49 (Figure 3B). Besides the inter-strand hydrogen bonds, the hydrophobic residues located at the interface (Phe39, Thr40, and Phe51 – Figure 3C) exhibit a decrease of their surface accessible solvent area (SASA) after dimerization. The SASA reductions of hydrophobic residues favour the binding of the monomers. The occurrence of π-π interactions between three aromatic pairs (Tyr52-Tyr52, Phe39-Phe51, and Phe51-Phe51) were also observed along the simulations. Therefore, the dimerization is favoured by hydrogen bonds, hydrophobic interactions, and π-π interactions.

**Figure 3:**
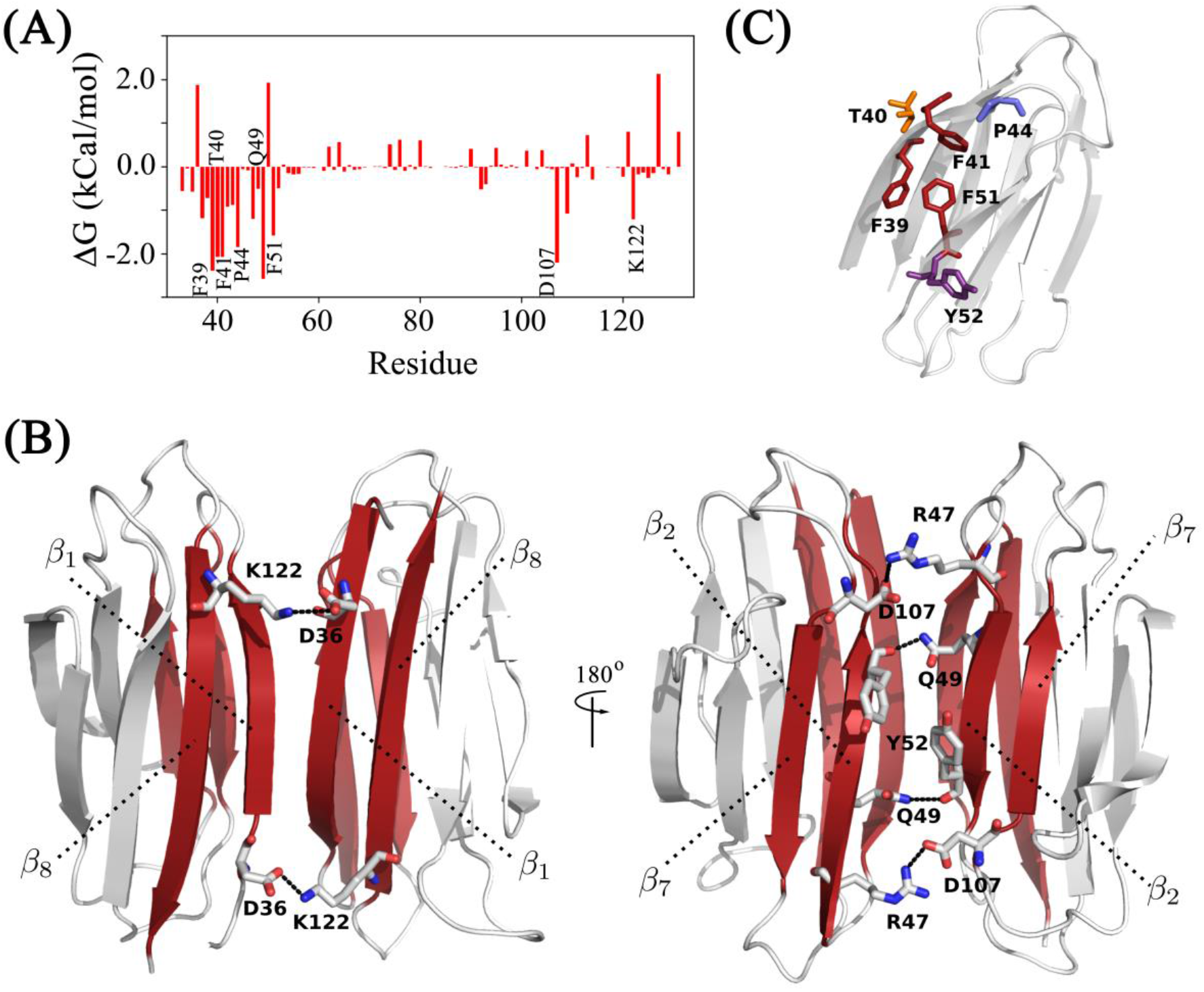
Analysis of the TMED1 GOLD dimerization properties. (A) Per-residue contribution to the total dimerization free energy of TMED1 dimer. Residues with ΔG < −1.5 kcal/mol were labelled. (B) Representation of the pairs of intermolecular hydrogen bonds (highlighted in black dotted lines) that contribute for the dimer stabilization. (C) Position of the hydrophobic and aromatic residues on the binding interface of the TMED1 GOLD domain. For simplicity, only one monomer was shown. The figure was prepared using Adobe Fireworks^®^ CS6, Origin® 8.5 and VMD [67].

Interestingly, most of those residues observed to be important for the TMED1 GOLD dimerization are conserved inside the γ subfamily in *Homo sapiens*. Thr40, Phe41, Pro44, Phe51, and Lys122 are highly conserved except in TMED6, the most different representative from the γ subfamily (Figure S3). Phe39, Tyr52, and Asp107 are conserved only in TMED1 and TMED5 (Figure S3). It was shown before that a chimeric TMED5 carrying the GOLD domain of TMED1 (instead of its own) can keep the transport of glycosylphosphatidylinositol-anchored proteins [66]. Therefore, besides their higher sequence identity (Figure 1B) and possibly a functional redundancy, their GOLD domains can indeed dimerize using similar strategies. The absence of some of the residues important for the dimerization of TMED1 GOLD might explain the reason why TMED7 is present in an equilibrium of monomer/dimer, and TMED3 is predominantly a monomer *in vivo* [24]. We could not find information about the TMED6 oligomerization in the literature.

Since the hydrogen bonds observed in the dimeric TMED1 GOLD structure have significant contributions to the free energy of interaction, we analysed the molecular dimension of this protein as a function of the ionic strength (Figure 4A). The protein’s elution profile shifted from a peak centred at 13.5 mL, corresponding to the monomer at high salt concentration, as confirmed by the SEC-MALS data (Figure 4A), towards lower elution volumes with the decrease in the salt concentration (Figure 4A). This means that the hydrodynamic radius associated with the protein structural organization increases with NaCl reduction, which we associate with a dimer formation (Figure 4A). It is important to emphasize that the shift in the elution volume observed in the SEC data for the TMED1 GOLD at low salt cannot be associated with a non-specific interaction with the Superdex75 matrix. Whether this was the case, it would be expected a higher protein retention into the column, leading to a greater elution volume than that observed with high salt concentration. In order to also test the possibility of an early elution based on an electrostatic repulsion between the protein and the column matrix, the solution pH at a fixed NaCl concentration of 20 mM was varied (Figure S4). There is no significant shift compared with the controlled profile associated with the dimeric structure (Figure S4). The same conclusion is reached when the buffering agent is changed, or acetonitrile is added to disturb hydrophobic association (Figure S4). The data give strength to the conclusion that the early elution volume observed at low NaCl concentrations is indeed due to a dimeric organization.

**Figure 4:**
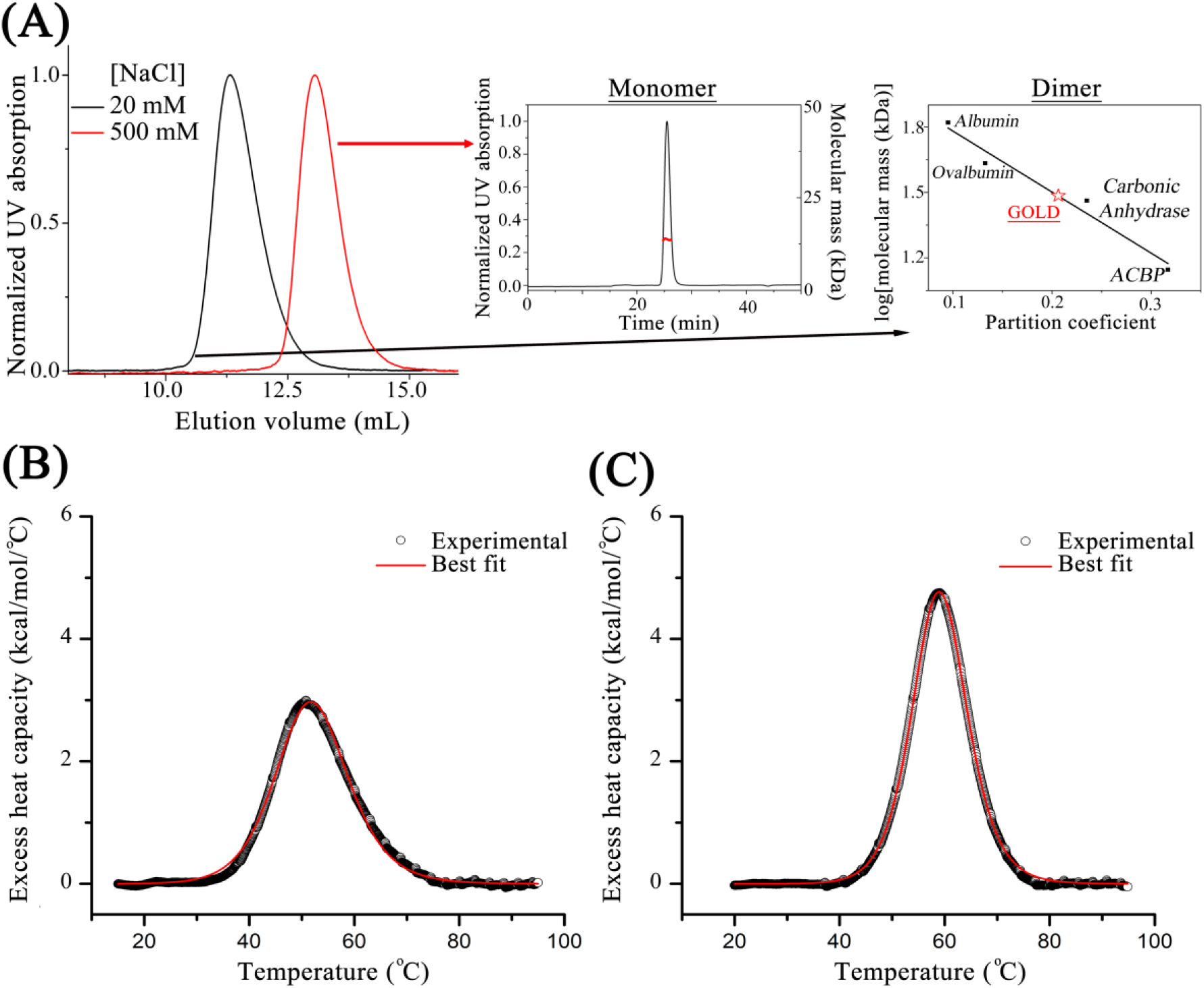
(A) SEC results of the TMED1 GOLD domain oligomerization propensity according to the salt concentration in solution. The GOLD domain oligomerization was analysed by SEC-MALS in 50 mM Tris/HCl, pH 8.0, 500 mM NaCl and by comparing the partition coefficient with that of standard proteins in 20 mM Tris/HCl, pH 8.0, 20 mM NaCl condition. (B) and (C) are the excess heat capacity of TMED1 GOLD at low (B) and high (C) ionic strengths. The baseline-corrected thermograms were deconvoluted with a non-two-state (B) or a two-state (C) model for protein unfolding. Experimental data are represented by black open circles, and the best fits are the solid red lines. The best-fit parameters were the following: (B) *T*_m_ = 50.9 ± 0.1°C, Δ*H*_cal_ = 52.6 ± 2.3 kcal/mol, Δ*H*_*vH*_ = 45.5 ± 0.3 kcal/mol, Δ*T*_*1/2*_ = 16.7 ± 0.1°C, Δ*H*_*vH*_/Δ*H*_*cal*_ = 0.87 ± 0.04; and (C) *T*_m_ = 58.8 ± 0.2°C, Δ*H*_cal_ = 64.1 ± 1.4 kcal/mol, Δ*H*_*vH*_ = 63.4 ± 0.6 kcal/mol, Δ*T*_*1/2*_ = 12.5 ± 0.1°C, Δ*H*_*vH*_/Δ*H*_*cal*_ = 0.99 ± 0.03.

We also investigated the effect of the ionic strength on the protein stability by DSC. Figures 4B and 4C illustrate the temperature dependence of the excess heat capacity of TMED1 GOLD at low and high NaCl concentrations, respectively. Addition of NaCl promotes protein structural stabilization: the denaturation of TMED1 GOLD is more thermally (ΔT_m_ = + 7.9°C) and enthalpically (+ 10.3 kcal/mol) stable as well as more cooperative (smaller linewidth at half height, ΔT_1/2_) at high ionic strength. In both conditions, the protein has shown an averaged thermal reversibility ratio of about 84% after each scan (Figure S5). Additionally, the DSC thermogram obtained at low ionic strength, corresponding to the dimer state, was asymmetric towards high temperatures, suggesting that the protein denaturation does not follow a standard reversible two-state process. Indeed, deconvolution of the thermogram indicated that the ratio between the van’t Hoff enthalpy change (ΔH_vH_) and the calorimetric enthalpy change (ΔH_cal_) is < 1, suggesting that the unfolding process of the TMED1 GOLD dimer likely involves the formation of intermediate states [68,69]. On the other hand, the ΔH_vH_/ΔH_cal_ ∼ 1 obtained at 300 mM NaCl indicates that the monomeric TMED1 GOLD unfolds following an ideal reversible two-state process, as shown for a variety of small single-domain globular proteins [69,70].

To further explore the unfolding mechanism of the TMED1 GOLD dimer, we used the web-based CalFitter software to test different protein unfolding models that could putatively explain the TMED1 GOLD temperature denaturation process obtained at low ionic strength. We performed a global data analysis using the same unfolding model to fit a set of thermal denaturation data obtained by both DSC and CD. The unfolding pathway that better simulated the combined experimental data was a model that involves a transition with a reversible step in equilibrium (native ↔ intermediate) and an irreversible step (intermediate → denatured), thus further suggesting the existence of an intermediate state (Figure S5C-E).

Taken together, our results indicate that the GOLD domain of TMED1 form dimers in solution in a salt-dependent way and that the dimer observed in the crystal structure is stable and energetically favourable, therefore likely being the *in vivo* organization of the domain.

### 3.4 Implications of the GOLD dimer structure and a model for TMED membrane anchoring

To expand the discussion about the orientation of the GOLD relative to the membrane surface, we calculated the electrostatic potential surface of both TMED1 and TMED5 GOLD dimer structures. Interestingly, we found that the dimers of TMED1 and TMED5 GOLD have a conserved feature absent in the TMED2 and TMED10 GOLD: a large negatively charged surface formed in the convex side (Figure 5A). This result suggests that this feature might be exclusive for the γ-subfamily. This characteristic gives strength to the proposed model regarding the orientation of the GOLD domains relative to the membrane surface, since both the Golgi and the ER membranes are enriched in anionic lipids, including phosphatidylinositol and sphingolipids [71]. Therefore, we hypothesize that the negatively charged surface of TMED1 GOLD remains exposed to the cytosol during their traffic in the early secretory pathway, while the concave side would face the membrane surface (Figure 5B). This orientation has been previously suggested for TMED5, although the authors based their conclusions only on a possible continuation of the C-terminal region [23]. Here we propose that the GOLD orientation is driven by the electrostatic repulsion between the membrane surface and the convex side of TMEDs 1 and 5, which is absent in TMEDs 2 and 10 GOLD. How this could play a role during the TMEDs-associated processes, especially in protein traffic, remains unknown. Future studies including the binding partners of these GOLD domains may provide some clues on the cargo-signal regions that drive them to the TMED-dependent COP insertion.

**Figure 5:**
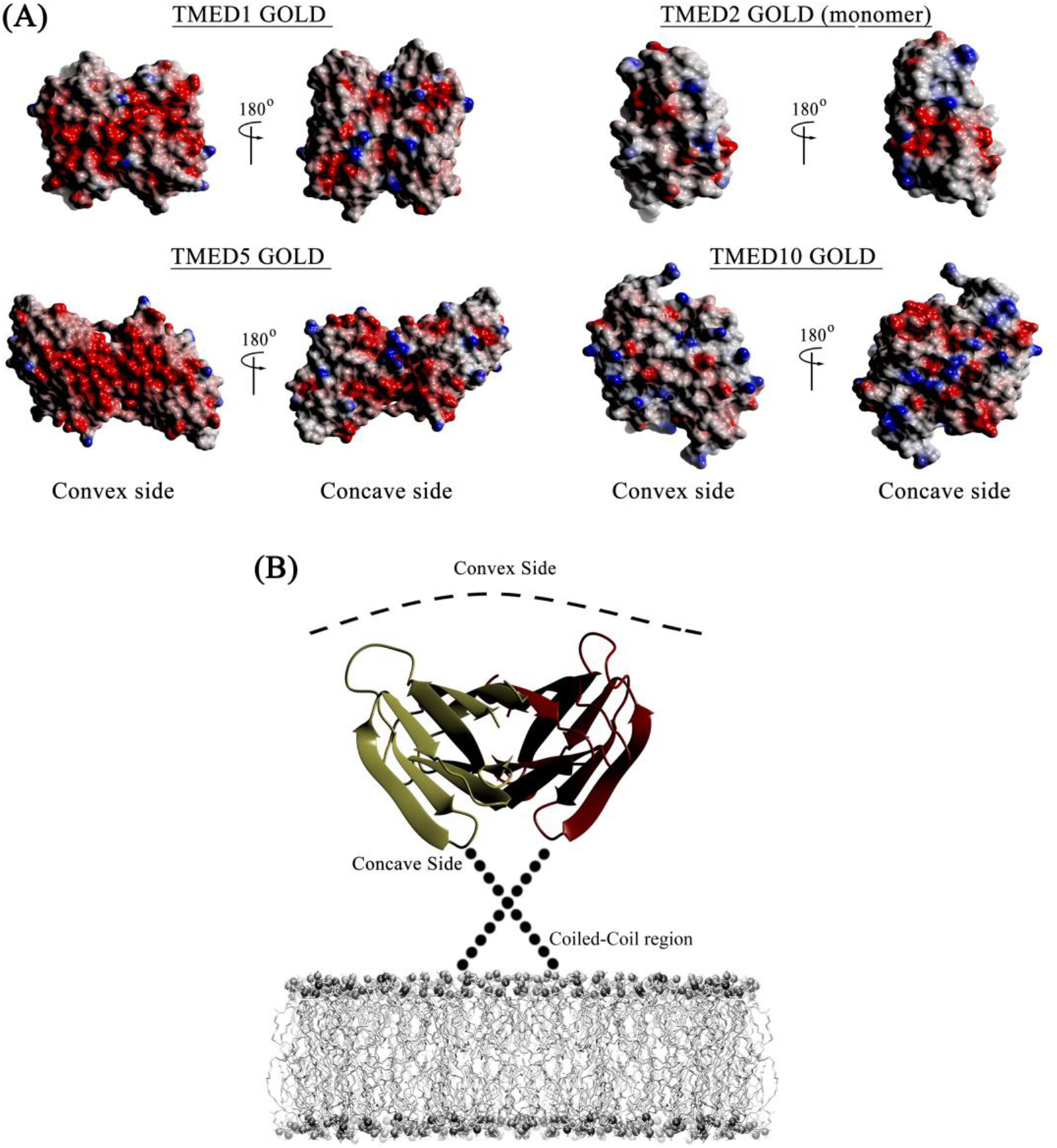
Proposed TMED1 dimerization model in subcompartments of the early secretory pathway. (A) Electrostatic surface potential maps of TMED1, TMED2 (PDB ID 5AZW - monomer), TMED5 (PDB ID 5GU5), and TMED10 (PDB ID 5AZY) GOLD structures, calculated using the Poisson-Boltzmann solver and default parameters for colouring within CCP4MG. The colour of the surface represents the electrostatic potential on the protein surface, going from blue (potential of +0.5 V) to red (potential of −0.5 V). (B) Proposed model for TMED1 orientation in the membranes, exposing the negatively charged surface formed by the convex side to the lumen. In this model, the concave side is oriented towards the membrane surface. The figure was prepared using Adobe Fireworks^®^ CS6 and CCP4MG.

Finally, it has been previously shown that TMED1 forms dimers in all subcompartments of the early secretory pathway, named ER, Golgi, and intermediate compartments [62]. This dimerization propensity was unique between different TMEDs [62]. Here, we showed that the TMED1 dimer formation is mediated by the isolated GOLD domain and future studies will be conducted to assess the participation and the relevance of the coiled-coil region on the TMED1 oligomerization.

## Acknowledgments

The authors thank the Brazilian agencies Conselho Nacional de Desenvolvimento Científico e Tecnológico (CNPq) and Fundação de Amparo à Pesquisa do Estado de São Paulo (FAPESP) for the financial support through Grants No. 2015/50366-7 and 306682/2018-4. DCAMM thanks the Conselho Nacional de Desenvolvimento Científico e Tecnológico (CNPq) for the MSc scholarship (134160/2018-5). LFSM acknowledges FAPESP for the post-doctoral Grant No. 2017/24669-8. The authors also thank the Molecular Biophysics Group at Sao Carlos Physics Institute of the University of Sao Paulo for allowing access to the DSC calorimeter and SEC MALS (FAPESP Grant number: 15/16812-0).

## Data availability

The dataset generated and analysed during the current study, together with the raw data, are available in the Protein Data Bank under the ID 7K49 and/or upon reasonable request to the correspondent authors. All data used in the discussions are included in the main manuscript and in the supplementary information.

## References

1 Bonifacino, J. S. & Glick, B. S. The mechanisms of vesicle budding and fusion. Cell 116, 153–166, doi:10.1016/s0092-8674(03)01079-1 (2004).

2 Lee, M. C. S., Miller, E. A., Goldberg, J., Orci, L. & Schekman, R. Bi-directional protein transport between the ER and Golgi. Annual Review of Cell and Developmental Biology 20, 87–123, doi:10.1146/annurev.cellbio.20.010403.105307 (2004).

3 Dancourt, J. & Barlowe, C. Protein Sorting Receptors in the Early Secretory Pathway. Annual Review of Biochemistry, Vol 79 79, 777–802, doi:10.1146/annurev-biochem-061608-091319 (2010).

4 Gomez-Navarro, N. & Miller, E. Protein sorting at the ER-Golgi interface. Journal of Cell Biology 215, 769–778, doi:10.1083/jcb.201610031 (2016).

5 Barlowe, C. et al.. COPII - A MEMBRANE COAT FORMED BY SEC PROTEINS THAT DRIVE VESICLE BUDDING FROM THE ENDOPLASMIC-RETICULUM. Cell 77, 895–907, doi:10.1016/0092-8674(94)90138-4 (1994).

6 Malhotra, V., Serafini, T., Orci, L., Shepherd, J. C. & Rothman, J. E. PURIFICATION OF A NOVEL CLASS OF COATED VESICLES MEDIATING BIOSYNTHETIC PROTEIN-TRANSPORT THROUGH THE GOLGI STACK. Cell 58, 329–336, doi:10.1016/0092-8674(89)90847-7 (1989).

7 Kuehn, M. J., Herrmann, J. M. & Schekman, R. COPII-cargo interactions direct protein sorting into ER-derived transport vesicles. Nature 391, 187–190 (1998).

8 Benyair, R., Ron, E. & Lederkremer, G. Z. PROTEIN QUALITY CONTROL, RETENTION, AND DEGRADATION AT THE ENDOPLASMIC RETICULUM. International Review of Cell and Molecular Biology, Vol 292 292, 197–280, doi:10.1016/b978-0-12-386033-0.00005-0 (2011).

9 Uhlen, M. et al.. The human secretome. Science Signaling 12, doi:10.1126/scisignal.aaz0274 (2019).

10 Strating, J. & Martens, G. J. M. The p24 family and selective transport processes at the ER-Golgi interface. Biology of the Cell 101, 495–509, doi:10.1042/bc20080233 (2009).

11 Carney, G. E. & Bowen, N. J. p24 proteins, intracellular trafficking, and behavior: Drosophila melanogaster provides insights and opportunities. Biology of the Cell 96, 271–278, doi:10.1016/j.biolcel.2004.01.004 (2004).

12 Pastor-Cantizano, N., Montesinos, J. C., Bernat-Silvestre, C., Marcote, M. J. & Aniento, F. p24 family proteins: key players in the regulation of trafficking along the secretory pathway. Protoplasma 253, 967–985, doi:10.1007/s00709-015-0858-6 (2016).

13 Vetrivel, K. S. et al. Dual roles of the transmembrane protein p23/TMP21 in the modulation of amyloid precursor protein metabolism. Molecular Neurodegeneration 2, doi:10.1186/1750-1326-2-4 (2007).

14 Yang, X. Y. et al. Identification of differentially expressed genes in metastatic and non-metastatic nasopharyngeal carcinoma cells by suppression subtractive hybridization. Cellular Oncology 27, 215–223 (2005).

15 Zheng, H. et al. TMED3 promotes hepatocellular carcinoma progression via IL-11/STAT3 signaling. Scientific Reports 6, doi:10.1038/srep37070 (2016).

16 Wang, X. C. et al. Transmembrane Emp24 Protein Transport Domain 6 is Selectively Expressed in Pancreatic Islets and Implicated in Insulin Secretion and Diabetes. Pancreas 41, 10–14, doi:10.1097/MPA.0b013e318223c7e4 (2012).

17 Saleem, S. et al. Drosophila melanogaster p24 trafficking proteins have vital roles in development and reproduction. Mechanisms of Development 129, 177–191, doi:10.1016/j.mod.2012.04.002 (2012).

18 Stamnes, M. A. et al. AN INTEGRAL MEMBRANE COMPONENT OF COATOMER-COATED TRANSPORT VESICLES DEFINES A FAMILY OF PROTEINS INVOLVED IN BUDDING. Proceedings of the National Academy of Sciences of the United States of America 92, 8011–8015, doi:10.1073/pnas.92.17.8011 (1995).

19 Denzel, A. et al. The p24 family member p23 is required for early embryonic development. Current Biology 10, 55–58, doi:10.1016/s0960-9822(99)00266-3 (2000).

20 Strating, J., van Bakel, N. H. M., Leunissen, J. A. M. & Martens, G. J. M. A Comprehensive Overview of the Vertebrate p24 Family: Identification of a Novel Tissue-Specifically Expressed Member. Molecular Biology and Evolution 26, 1707–1714, doi:10.1093/molbev/msp099 (2009).

21 Dominguez, M. et al. gp25L/emp24/p24 protein family members of the cis-Golgi network bind both COP I and II coatomer. Journal of Cell Biology 140, 751–765, doi:10.1083/jcb.140.4.751 (1998).

22 Nagae, M. et al. 3D Structure and Interaction of p24 beta and p24 delta Golgi Dynamics Domains: Implication for p24 Complex Formation and Cargo Transport. Journal of Molecular Biology 428, 4087–4099, doi:10.1016/j.jmb.2016.08.023 (2016).

23 Nagae, M. et al. Crystallographic analysis of murine p24 gamma 2 Golgi dynamics domain. Proteins-Structure Function and Bioinformatics 85, 764–770, doi:10.1002/prot.25242 (2017).

24 Connolly, D. J., O’Neill, L. A. J. & McGettrick, A. F. The GOLD Domain-containing Protein TMED1 Is Involved in Interleukin-33 Signaling. Journal of Biological Chemistry 288, 5616–5623, doi:10.1074/jbc.M112.403899 (2013).

25 Alves, J. C. et al. Interleukin-33 attenuates sepsis by enhancing neutrophil influx to the site of infection. Nature Medicine 16, 708–U113, doi:10.1038/nm.2156 (2010).

26 Schmitz, J. et al. IL-33, an interleukin-1-like cytokine that signals via the IL-1 receptor-related protein ST2 and induces T helper type 2-associated cytokines. Immunity 23, 479–490, doi:10.1016/j.immuni.2005.09.015 (2005).

27 Ohno, T., Morita, H., Arae, K., Matsumoto, K. & Nakae, S. Interleukin-33 in allergy. Allergy 67, 1203–1214, doi:10.1111/all.12004 (2012).

28 Liew, F. Y., Pitman, N. I. & McInnes, I. B. Disease-associated functions of IL-33: the new kid in the IL-1 family. Nature Reviews Immunology 10, 103–110, doi:10.1038/nri2692 (2010).

29 Doyle, S. L. et al. The GOLD domain-containing protein TMED7 inhibits TLR4 signalling from the endosome upon LPS stimulation. Nature Communications 3, doi:10.1038/ncomms1706 (2012).

30 Micsonai, A. et al. BeStSel: a web server for accurate protein secondary structure prediction and fold recognition from the circular dichroism spectra. Nucleic Acids Research 46, W315–W322, doi:10.1093/nar/gky497 (2018).

31 Micheletto, M. C., Mendes, L. F. S., Basso, L. G. M., Fonseca-Maldonado, R. G. & Costa-Filho, A. J. Lipid membranes and acyl-CoA esters promote opposing effects on acyl-CoA binding protein structure and stability. International Journal of. Biological Macromolecules 102, 284–293, doi:10.1016/j.ijbiomac.2017.03.197 (2017).

32 Grek, S. B., Davis, J. K. & Blaber, M. An efficient, flexible-model program for the analysis of differential scanning calorimetry protein denaturation data. Protein and Peptide Letters 8, 429–436, doi:10.2174/0929866013409184 (2001).

33 Mazurenko, S. et al. CalFitter: a web server for analysis of protein thermal denaturation data. Nucleic Acids Research 46, W344–W349, doi:10.1093/nar/gky358 (2018).

34 Kabsch, W. XDS. Acta Crystallographica Section D-Biological Crystallography 66, 125–132, doi:10.1107/s0907444909047337 (2010).

35 Winn, M. D. et al. Overview of the CCP4 suite and current developments. Acta Crystallographica Section D-Structural Biology 67, 235–242, doi:10.1107/s0907444910045749 (2011).

36 Vagin, A. & Teplyakov, A. Molecular replacement with MOLREP. Acta Crystallographica Section D-Biological Crystallography 66, 22–25, doi:10.1107/s0907444909042589 (2010).

37 Cowtan, K. The Buccaneer software for automated model building. 1. Tracing protein chains. Acta Crystallographica Section D-Biological Crystallography 62, 1002–1011, doi:10.1107/s0907444906022116 (2006).

38 Murshudov, G. N. et al. REFMAC5 for the refinement of macromolecular crystal structures. Acta Crystallographica Section D-Structural Biology 67, 355–367, doi:10.1107/s0907444911001314 (2011).

39 Emsley, P., Lohkamp, B., Scott, W. G. & Cowtan, K. Features and development of Coot. Acta Crystallographica Section D-Biological Crystallography 66, 486–501, doi:10.1107/s0907444910007493 (2010).

40 Chen, V. B. et al. MolProbity: all-atom structure validation for macromolecular crystallography. Acta Crystallographica Section D-Structural Biology 66, 12–21, doi:10.1107/s0907444909042073 (2010).

41 McNicholas, S., Potterton, E., Wilson, K. S. & Noble, M. E. M. Presenting your structures: the CCP4mg molecular-graphics software. Acta Crystallographica Section D-Structural Biology 67, 386–394, doi:10.1107/s0907444911007281 (2011).

42 Armenteros, J. J. A. et al. SignalP 5.0 improves signal peptide predictions using deep neural networks. Nature Biotechnology 37, 420-+, doi:10.1038/s41587-019-0036-z (2019).

43 Jones, D. T., Taylor, W. R. & Thornton, J. M. THE RAPID GENERATION OF MUTATION DATA MATRICES FROM PROTEIN SEQUENCES. Computer Applications in the Biosciences 8, 275–282, doi:10.1093/bioinformatics/8.3.275 (1992).

44 Kumar, S., Stecher, G., Li, M., Knyaz, C. & Tamura, K. MEGA X: Molecular Evolutionary Genetics Analysis across Computing Platforms. Molecular Biology and Evolution 35, 1547–1549, doi:10.1093/molbev/msy096 (2018).

45 Madeira, F. et al. The EMBL-EBI search and sequence analysis tools APIs in 2019. Nucleic Acids Research 47, W636–W641, doi:10.1093/nar/gkz268 (2019).

46 Martinez, L., Andrade, R., Birgin, E. G. & Martinez, J. M. PACKMOL: A Package for Building Initial Configurations for Molecular Dynamics Simulations. Journal of Computational Chemistry 30, 2157–2164, doi:10.1002/jcc.21224 (2009).

47 Martinez, J. M. & Martinez, L. Packing optimization for automated generation of complex system’s initial configurations for molecular dynamics and docking. Journal of Computational Chemistry 24, 819–825, doi:10.1002/jcc.10216 (2003).

48 Phillips, J. C. et al. Scalable molecular dynamics with NAMD. Journal of Computational Chemistry 26, 1781–1802, doi:10.1002/jcc.20289 (2005).

49 MacKerell, A. D. et al. All-atom empirical potential for molecular modeling and dynamics studies of proteins. Journal of Physical Chemistry B 102, 3586–3616, doi:10.1021/jp973084f (1998).

50 Price, D. J. & Brooks, C. L. A modified TIP3P water potential for simulation with Ewald summation. Journal of Chemical Physics 121, 10096–10103, doi:10.1063/1.1808117 (2004).

51 Darden, T., York, D. & Pedersen, L. PARTICLE MESH EWALD -AN N.LOG(N) METHOD FOR EWALD SUMS IN LARGE SYSTEMS. Journal of Chemical Physics 98, 10089–10092, doi:10.1063/1.464397 (1993).

52 Miyamoto, S. & Kollman, P. A. SETTLE - AN ANALYTICAL VERSION OF THE SHAKE AND RATTLE ALGORITHM FOR RIGID WATER MODELS. Journal of Computational Chemistry 13, 952–962, doi:10.1002/jcc.540130805 (1992).

53 Chen, J., Sawyer, N. & Regan, L. Proteinprotein interactions: General trends in the relationship between binding affinity and interfacial buried surface area. Protein Science 22, 510–515, doi:10.1002/pro.2230 (2013).

54 Lee, B. & Richards, F. M. INTERPRETATION OF PROTEIN STRUCTURES - ESTIMATION OF STATIC ACCESSIBILITY. Journal of Molecular Biology 55, 379-&, doi:10.1016/0022-2836(71)90324-x (1971).

55 Swanson, J. M. J., Henchman, R. H. & McCammon, J. A. Revisiting free energy calculations: A theoretical connection to MM/PBSA and direct calculation of the association free energy. Biophysical Journal 86, 67–74, doi:10.1016/s0006-3495(04)74084-9 (2004).

56 Kollman, P. A. et al. Calculating structures and free energies of complex molecules: Combining molecular mechanics and continuum models. Accounts of Chemical Research 33, 889–897, doi:10.1021/ar000033j (2000).

57 Lee, T.-S. et al. GPU-Accelerated Molecular Dynamics and Free Energy Methods in Amber18: Performance Enhancements and New Features. Journal of Chemical Information and Modeling 58, 2043–2050, doi:10.1021/acs.jcim.8b00462 (2018).

58 Case, D. A. et al. The Amber biomolecular simulation programs. Journal of Computational Chemistry 26, 1668–1688, doi:10.1002/jcc.20290 (2005).

59 Miller, B. R., III et al. MMPBSA.py: An Efficient Program for End-State Free Energy Calculations. Journal of Chemical Theory and Computation 8, 3314–3321, doi:10.1021/ct300418h (2012).

60 Contreras, I., Yang, Y. D., Robinson, D. G. & Aniento, F. Sorting signals in the cytosolic tail of plant p24 proteins involved in the interaction with the COPII coat. Plant and Cell Physiology 45, 1779–1786, doi:10.1093/pcp/pch200 (2004).

61 Marzioch, M. et al. Erp1p and Erp2p, partners for Emp24p and Erv25p in a yeast p24 complex. Molecular Biology of the Cell 10, 1923–1938, doi:10.1091/mbc.10.6.1923 (1999).

62 Jenne, N., Frey, K., Brugger, B. & Wieland, F. T. Oligomeric state and stoichiometry of p24 proteins in the early secretory pathway. Journal of Biological Chemistry 277, 46504–46511, doi:10.1074/jbc.M206989200 (2002).

63 Emery, G., Rojo, M. & Gruenberg, J. Coupled transport of p24 family members. Journal of Cell Science 113, 2507–2516 (2000).

64 Ciufo, L. F. & Boyd, A. Identification of a lumenal sequence specifying the assembly of Emp24p into p24 complexes in the yeast secretory pathway. Journal of Biological Chemistry 275, 8382–8388, doi:10.1074/jbc.275.12.8382 (2000).

65 Campanera, J. M. & Pouplana, R. MMPBSA Decomposition of the Binding Energy throughout a Molecular Dynamics Simulation of Amyloid-Beta (A beta(10-35)) Aggregation. Molecules 15, 2730–2748, doi:10.3390/molecules15042730 (2010).

66 Theiler, R. et al. The alpha-Helical Region in p24 gamma(2) Subunit of p24 Protein Cargo Receptor Is Pivotal for the Recognition and Transport of Glycosylphosphatidylinositol-anchored Proteins. Journal of Biological Chemistry 289, 16835–16843, doi:10.1074/jbc.M114.568311 (2014).

67 Humphrey, W., Dalke, A. & Schulten, K. VMD: Visual molecular dynamics. Journal of Molecular Graphics & Modelling 14, 33–38, doi:10.1016/0263-7855(96)00018-5 (1996).

68 Privalov, P. L. & Potekhin, S. A. Scanning microcalorimetry in studying temperature-induced changes in proteins. Methods Enzymol 131, 4–51, doi:10.1016/0076-6879(86)31033-4 (1986).

69 Freire, E. Differential scanning calorimetry. Methods Mol Biol 40, 191–218, doi:10.1385/0-89603-301-5:191 (1995).

70 Privalov, P. L. Stability of proteins: small globular proteins. Adv Protein Chem 33, 167–241, doi:10.1016/s0065-3233(08)60460-x (1979).

71 van Meer, G. & de Kroon, A. I. P. M. Lipid map of the mammalian cell. Journal of Cell Science 124, 5–8, doi:10.1242/jcs.071233 (2011).

