## SUPPLEMENTARY INFORMATION for "Structural and thermodynamic analyses of human TMED1 (p24γ1) Golgi dynamics"

[1] Laboratório de Biofísica Molecular, Faculdade de Filosofia, Ciências e Letras de Ribeirão Preto, Universidade de São Paulo, Ribeirão Preto, SP, Brasil

[2] Laboratório de Cristalografia de Proteínas, Departamento de Física e Química, Faculdade de Ciências Farmacêuticas de Ribeirão Preto, Universidade de São Paulo, Ribeirão Preto, SP, Brazil.

[3] Laboratório de Ciências Físicas, Centro de Ciência e Tecnologia, Universidade Estadual do Norte Fluminense Darcy Ribeiro, Campos dos Goytacazes, RJ, Brasil.

**>TMED1 GOLD domain with the non-native aminoacids in the N-terminus**

MGSSHHHHHHSSGLVPRGSHMENLYFQGAGPPPIQDGEFTFLLPAGRKQC
FYQSAPANASLETEYQVIGGAGLDVDFTLESPQGVLLVSESRKADGVHTV
EPTEAGDYKLCFDNSFSTISEKLVFFELIFDSLQD

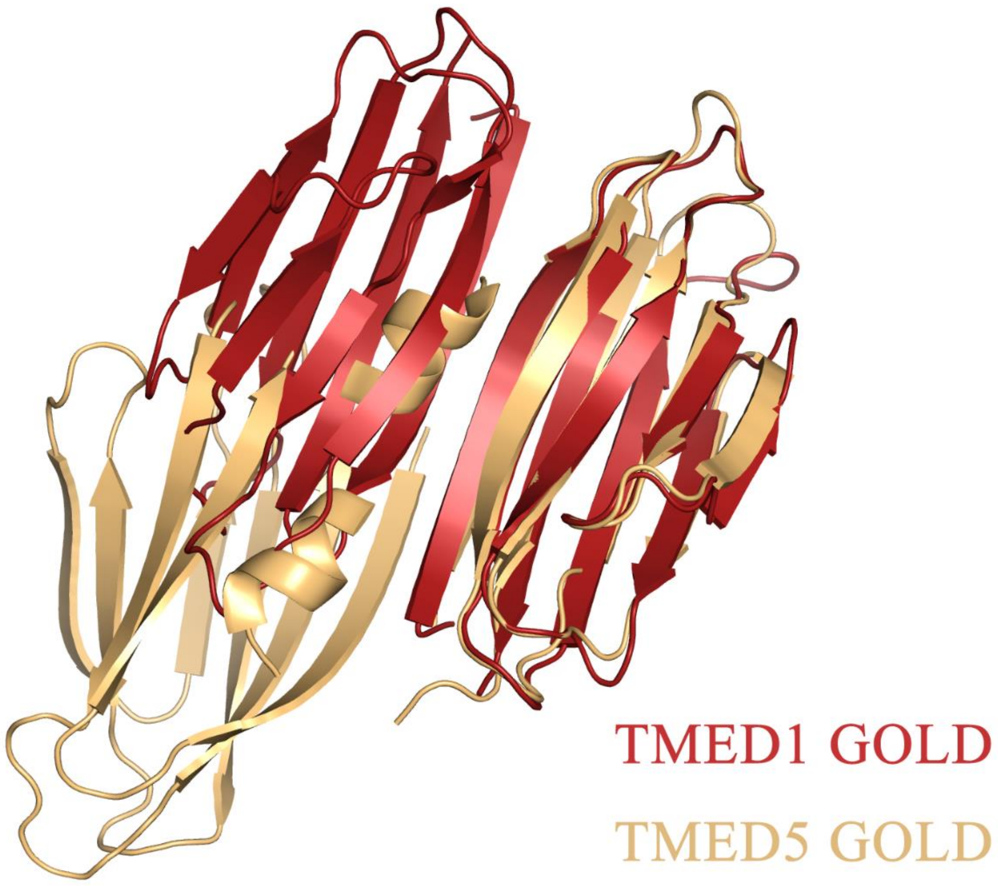


**Figure S1**: Structure superposition of the TMED1 GOLD (dark red) and TMED5 GOLD (gold) dimers using Pymol.

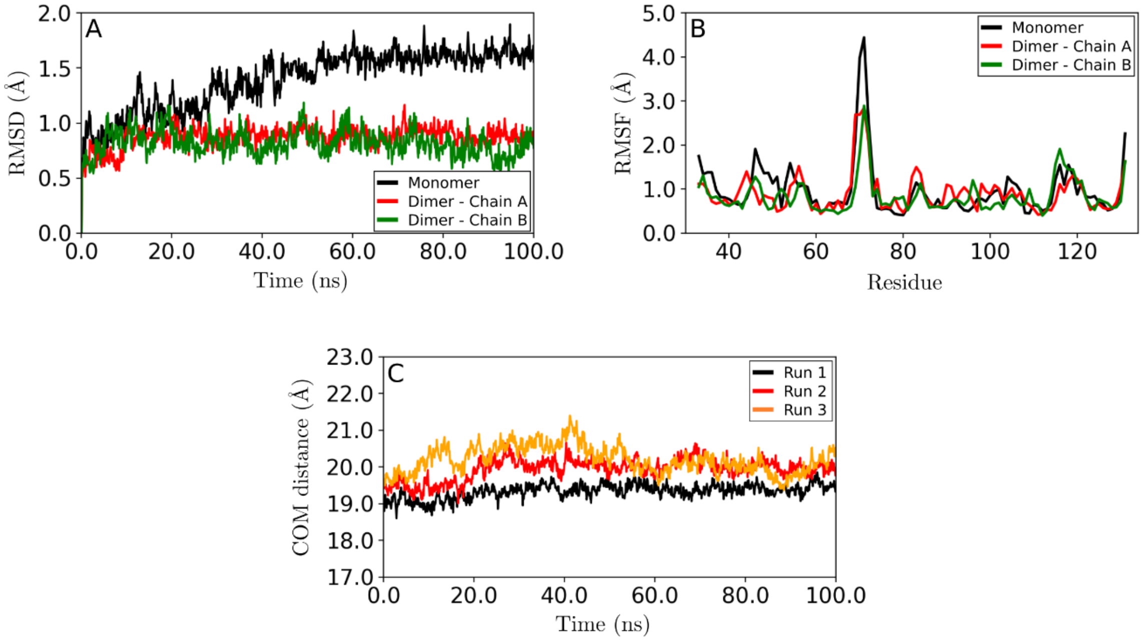


**Figure S2**: (A) RMSD along the trajectories for the backbone atoms of the monomer (black curve) and each monomer of the dimer (red and green curves). (B) Average Cα RMSD as a function of residue number. (C) Center of mass (COM) distance as function of time for each production simulation.

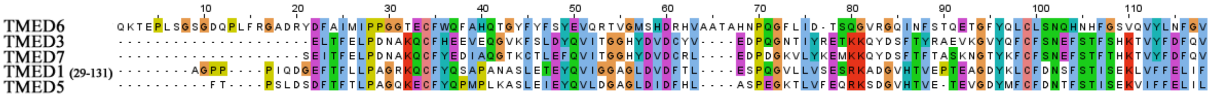


**Figure S3**. Multiple sequence alignment of the TMED GOLD representatives inside the γ subfamily in *Homo sapiens.* Sequences were aligned using Clustal Omega [1] and the figure, with the statistical analyses, was generated using Jalview (https://www.jalview.org/ accessed in 2020). There is a 28-residue shift between the numbered residues in the sequence alignment and the numbered residues in the PDB file. This is due to the non-native amino acids carried in the protein expression as a tag at the N-terminus.

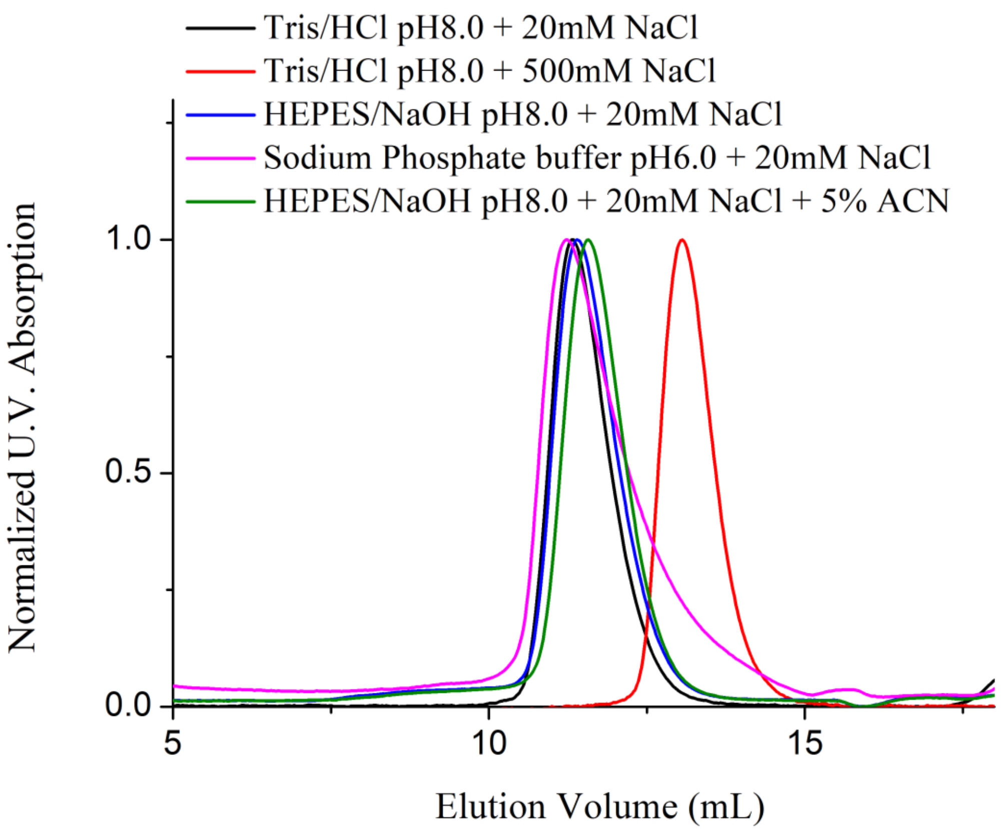


**Figure S4.** SEC results of the TMED1 GOLD domain used as control experiments. The data using 20 mM Tris/HCl + 20 or 500 mM NaCl are shown in the figure as examples of a dimeric and monomeric protein elution profiles. Changes in pH, buffering agent or addition of a polar solution does significantly change the profile associated with the dimeric peak. This is an indicative that the early elution of the associated-dimeric peak is not caused by electrostatic repulsion, unusual hydrophobic association or an unspecific effect caused by the buffering agent.

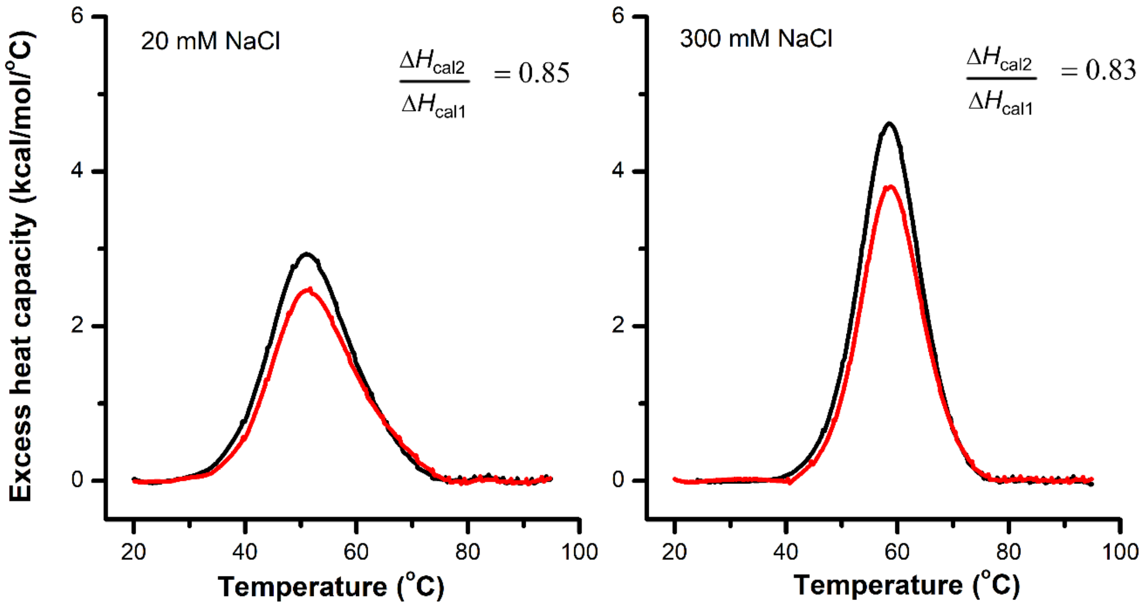


**Figure S5**. Thermal reversibility of TMED1 GOLD domain at low (left) and high (right) ionic strengths as assessed by DSC. The calorimetric enthalpy changes of the second heating scan (red) was about 83-85% of that of the first heating scan.

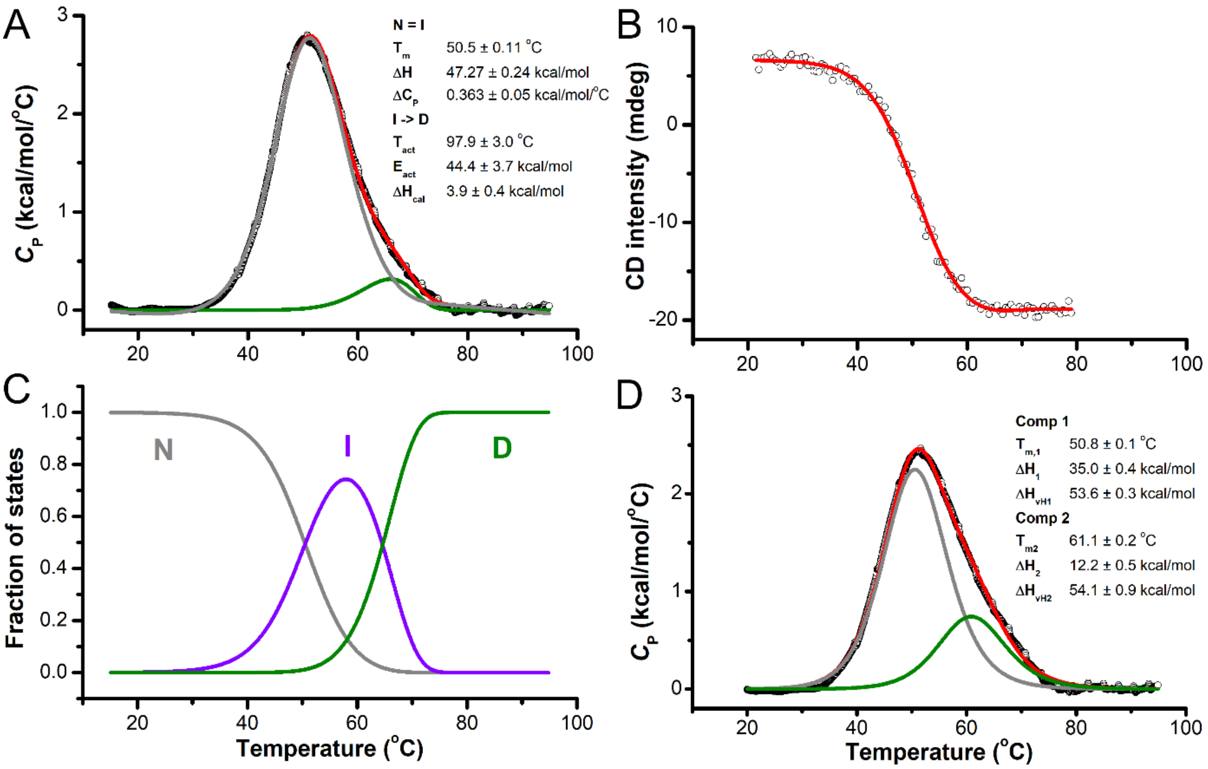


**Figure S6**. Analysis of the DSC and CD thermal denaturation data by CalFitter (A-C) and Microcal Origin (D) softwares. (A-C) CalFitter analysis. A cubic baseline were subtracted from the thermogram obtained at 20 mM NaCl and the resulting heat capacity profile along with the thermal denaturation of TMED1 GOLD, as monitored by CD (the ellipticity at 205 nm, panel B), were subjected to a global analysis using CalFitter web-server. The best-fit model of the protein denaturation was the N = I -> D, where N, I, and D stand for the native, intermediate, and denatured states, respectively. The "=" and "->" denote the equilibrium and irreversible steps, respectively. The best-fit parameters are shown in panel A and the temperature dependence of the fraction of each state is illustrated in panel C. For further information regarding the different protein unfolding models, please check Mazurenko *et al* [2]. (D) The thermogram in panel A was also simulated with a non-two-state process with two independent transitions using the Microcal Origin software. The best-fit parameters are shown in the plot. Experimental data are represented by black open circles, whereas colored solid lines denote the calculated curves.

### REFERENCES

1) Madeira F, Park YM, Lee J, Buso N, Gur T, Madhusoodanan N, Basutkar P, Tivey ARN, Potter SC, Finn RD, Lopez R. The EMBL-EBI search and sequence analysis tools APIs in 2019. Nucleic Acids Res. 2019 Jul 2;47(W1):W636-W641.

2) Mazurenko S, Stourac J, Kunka A, Nedeljkovic S, Bednar D, Prokop Z, Damborsky J. CalFitter: a web server for analysis of protein thermal denaturation data. Nucleic Acids Res. 2018 Jul 2;46(W1):W344-W349.
